# Spectral requirements for cooperation

**DOI:** 10.64898/2026.04.07.716994

**Authors:** Lior Pachter

## Abstract

We introduce a spectral existence criterion for the evolution of cooperation in the form of the inequality *λ*_max_*b > c*, where *λ*_max_ is the leading eigenvalue of an interaction operator encoding population structure, and *b* and *c* represent benefit and cost tradeoffs, respectively. Nowak’s five rules for the evolution of cooperation correspond to cases in which the cooperation condition reduces to a scalar assortment coefficient. These results follow from the Price equation, which sheds light on a long-standing debate on the role of inclusive fitness and evolutionary dynamics in explaining the evolution of cooperation.

## Introduction

In “Five rules for the evolution of cooperation” (Nowak, 2006), the author describes five distinct mechanisms for the evolution of cooperation and presents inequalities on parameters for each mechanism specifying when natural selection favors cooperative behavior. The mechanisms are kin selection, direct reciprocity, indirect reciprocity, network reciprocity, and group selection, and have been cited as characterizing all cooperation scenarios in nature (Nowak, 2012).

A natural question is how the five rules for the evolution of cooperation proposed in (Nowak, 2006) relate to the Price equation (Price, 1970), which was expressly introduced to study the evolutionary conditions underlying altruism (Harman, 2010). In (Nowak, 2006), the Price equation is mentioned only in a parenthetical remark (without formal citation) in a discussion of kin selection, despite the fact that the kin selection bound connecting relatedness to the conditions for cooperation is a direct consequence of the Price equation (Frank, 1997). Moreover, the kin selection cooperation bound is of the form *rb > c*, where *b* denotes the benefit to the recipient and *c* the cost to the actor (Hamilton, 1964), and while interpretations of *r* vary, this inequality appears to underlie all of the inequalities of (Nowak, 2006), as is evident in Table 1. This is no coincidence; it happens because the condition for cooperation reduces to the sign of a covariance between fitness and a cooperative trait, a point that is familiar to researchers in social evolution (Frank, 2012), but that is not formally established in (Nowak, 2006) or the papers associated with it (Nowak and May, 1992; Nowak and Sigmund, 1998; Traulsen and Nowak, 2006; Ohtsuki et al., 2006).

**Table 1.**
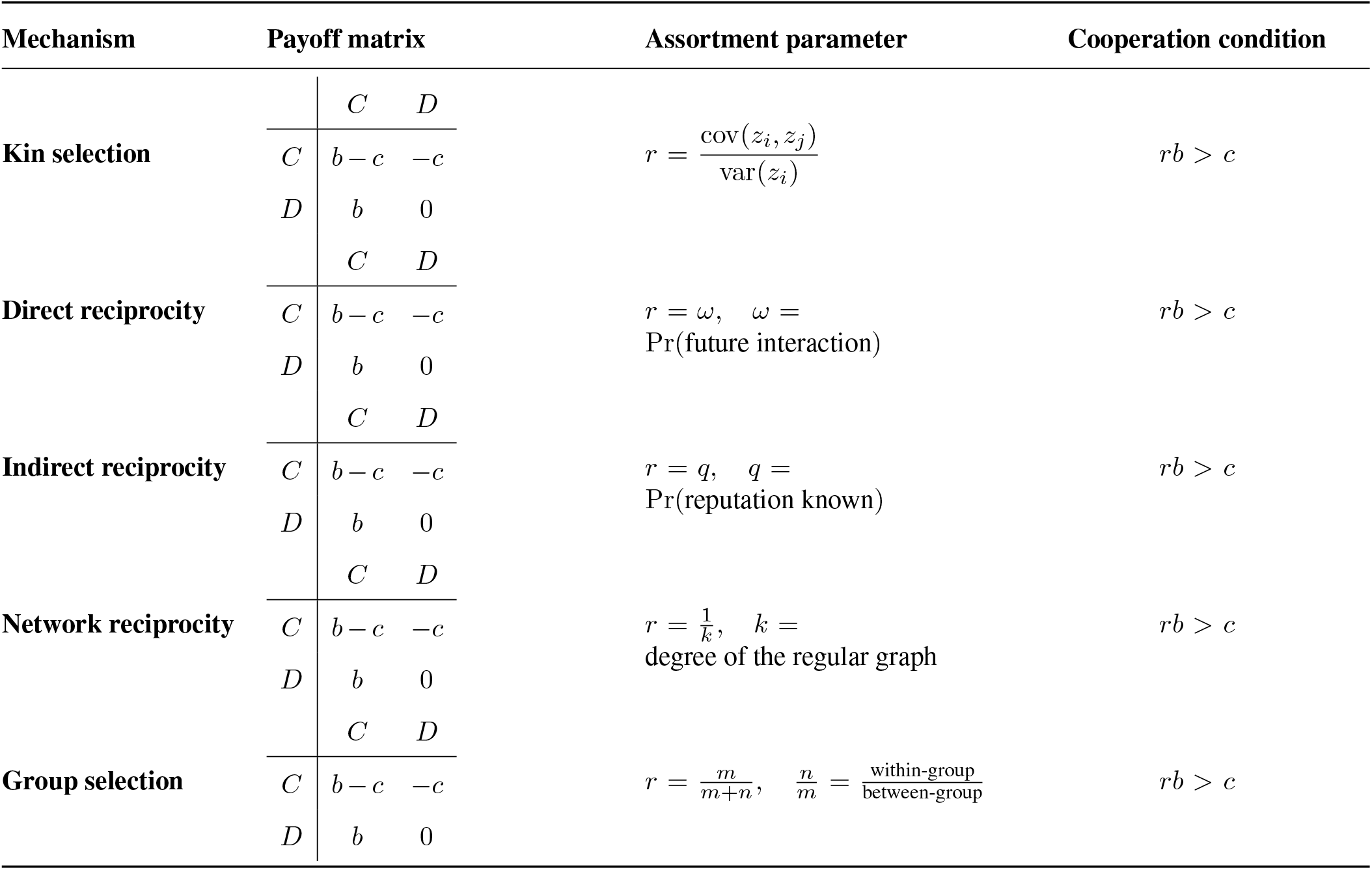
Five mechanisms for the evolution of cooperation described in (Nowak, 2006), expressed using a common donation-game payoff matrix. All mechanism-specific effects enter through the assortment parameter *r*, yielding the universal cooperation condition *rb > c*.

The discrete Price equation (Price, 1970) in the notation of (Felce et al., 2025) is stated as follows:

### Theorem 1

(Price (1970))

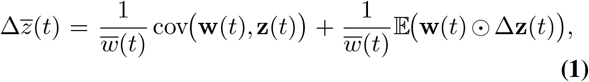

where **z**(*t*) = (*z*_1_(*t*),…, *z*_*n*_(*t*)) is the vector of trait values for *n* individuals or subpopulations at time *t*, **w**(*t*) = (*w*_1_(*t*),…, *w*_*n*_(*t*)) is the corresponding vector of Wrightian fitnesses, and expectations are computed with a weighting proportional to relative frequencies (*q*_1_(*t*),…, *q*_*n*_(*t*)) of population sizes (*p*_1_ (*t*),…, *p* (*t*)), i.e., 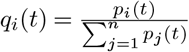. The study of cooperation inequalities is the application of the Price equation to the binary trait indicating cooperation or lack thereof. The first term on the right-hand side of Eq. (1) quantifies selection, while the second term represents transmission bias (e.g., mutation).

Nowak’s five rules are all direct corollaries of Eq. (1), requiring only an assumption of weak selection (Fisher, 1930; Haldane, 1932) that allows for linearization of fitness, and mechanism-specific assumptions about interaction structure (Kimura, 1964). Subsequently, each rule emerges from an assortment coefficient based on interaction structure between individuals in a population or between subpopulations. In all cases, the transmission term is suppressed, although we will see that if it is not, one obtains mutation-based cooperation rules. The specific forms of the assortment coefficients can be motivated by Moran processes (Moran, 1958), although these are not strictly required to derive the cooperation inequalities of (Nowak, 2006), or generalizations thereof.

The discrete Price equation has a continuum limit analog known as the continuous Price equation, which is derived by taking the limit as time differences converge to zero. The discrete Price equation then becomes a differential identity that relates the instantaneous change in the mean trait to selection and transmission via a change of coordinates from Wrightian fitness **w**(*t*) to Malthusian fitness **m**(*t*) (Felce et al., 2025):

### Theorem 2

(Price (1972); Ellner et al. (2011))

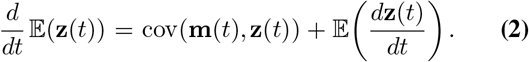

The existence of this continuum analog of Eq. (1) raises the question of whether Nowak’s five cooperation rules possess continuous counterparts. We show that the continuous Price equation can be used to derive conditions for evolvability via spectral inequalities involving eigenvalues of operators that encode the interaction architecture of the population.

### Selection

Nowak’s five rules for the evolution of cooperation can all be derived in three steps (Taylor and Frank, 1996):

1. Linearization of fitness under weak selection.
2. Derivation of an assortment coefficient.
3. Application of the Price equation.

The essence of the rules is step 3, namely the Price equation, and so the five rules are all corollaries of Theorem 1:

#### Corollary 1

(Hamilton (1964); Price (1970); Grafen (1985)) Each of Nowak’s five rules for the evolution of cooperation is a direct consequence of the Price equation.

*Proof:* An assumption of weak selection under the Prisoner’s Dilemma payoff matrix (Table 1) is tantamount to the assertion that fitness can be linearized as

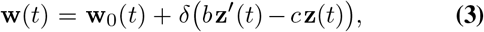

where **w**_0_(*t*) is trait independent, **z**(*t*) ∈ {0, 1} ^*n*^ is the vector of focal cooperative traits, **z**^′^(*t*) ∈ [0, 1]^*n*^ is the vector of expected cooperative traits of partners, and *δ* is small so that higher-order terms in *δ* can be ignored. Substituting Eq. (3) into the selection term of Eq. (1), noting that cov(**w**_0_(*t*), **z**(*t*)) = 0, and retaining only first-order terms in *δ* yields

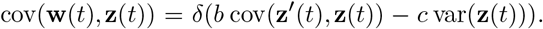

Define

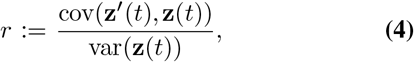

so that cov(**z**^′^(*t*), **z**(*t*)) = *r* var(**z**(*t*)). The quantity *r* is the least-squares regression coefficient (slope) of the partner trait **z**^′^(*t*) on the focal trait **z**(*t*) with respect to the Price weights, and may be interpreted as an assortment coefficient (Price, 1970; Grafen, 1985). Under weak selection, Eq. (4) is evaluated under the neutral reference process. At first order in the selection strength, the resulting regression coefficient *r* depends only on population structure and the interaction or updating mechanism and is therefore time-independent. Using *r*, the weak-selection assumption becomes

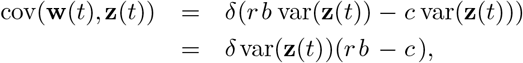

and the “five rules” arise because each mechanism in (Nowak, 2006) corresponds to a different parameterization of *r* applied to the same covariance identity. Specifically, since generically var(**z**(*t*)) *>* 0,

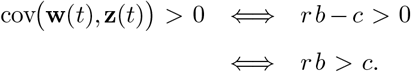

Thus, for any mechanism whose population structure enters only through *r*, and for which the transmission term in the Price equation can be ignored (faithful reproduction), cooperation is favored if, and only if, *rb > c*.

We now describe *r* for each of the five mechanisms in (Nowak, 2006), thus completing the proof of the corollary.

#### (i) Kin selection

Here *i* indexes individual ancestral lineages (1 ≤ *i* ≤ *n*), and the Price equation is applied at the individual level with uniform weights *q*_*i*_(*t*) = 1*/n*. In Nowak’s kin-selection model, assortment arises from genetic relatedness between donor and recipient. Let *r*_kin_ denote the regression coefficient of partner trait on focal trait, i.e.,

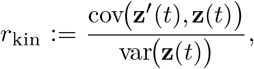

and let *r* = *r*_kin_. The Price equation condition *rb > c* therefore becomes *r*_kin_*b > c*, which is the rule for kin selection in (Nowak, 2006, Table 1).

#### (ii) Direct reciprocity

We apply the Price equation at the individual level, with *i* indexing individuals. In the direct reciprocity model, individuals play a repeated donation game (or repeated Prisoner’s Dilemma): after each interaction, the same pair continues with probability *ω* and terminates with probability 1 − *ω*. Thus, *ω* governs the temporal persistence of interaction partners.

Under weak selection, expectations in the Price equation may be evaluated under the neutral interaction process, since selection perturbs payoffs but not the interaction sequence at first order (Kimura, 1964). Let *z*_*i*_(*t*) ∈ {0, 1} denote the focal individual’s action (cooperate or defect) at time *t*, and let 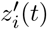 denote the partner’s action at time *t*. In direct reciprocity, assortment does not arise from population structure but from contingent behavior: a partner’s future action can depend on the focal individual’s current action.

In the simplest faithful-response idealization, when the same pair meets again the partner matches the focal individual’s previous action in expectation, whereas upon re-pairing the partner’s action is an independent draw from the population. Under this behavioral rule, the conditional expectation of the partner’s next action is

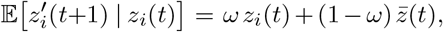

where 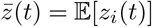 is the population mean action at time *t*. Because this dependence is linear in *z*_*i*_(*t*) with slope *ω*, the resulting Price-equation regression coefficient of the partner’s next action on the focal individual’s current action reduces to *ω* under this idealization (Grafen, 1985). Thus,

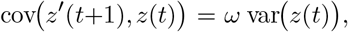

so the effective assortment coefficient for direct reciprocity is *r*_dir_ := *ω*. Substituting *r*_dir_ into the Price-equation condition recovers Nowak’s direct-reciprocity rule in this standard repeated-interaction setting.

#### (iii) Indirect reciprocity

Here *i* indexes individuals, and the Price equation is applied at the individual level. In the indirect reciprocity model, individuals engage in a reputation-based donation game: with probability *q* an individual observes the reputation of a prospective interaction partner, and with probability 1 − *q* the reputation is unobserved. Thus, *q* governs the availability of reputational information across interactions.

Under weak selection, expectations in the Price equation may be evaluated under the neutral reputation dynamics, since selective effects enter only at first order (Kimura, 1964). Let *z*_*i*_(*t*) ∈ { 0, 1} denote the cooperative action of the focal individual at time *t*, and let 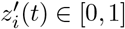 denote the probability that the interaction partner cooperates with *i* at time *t*. In indirect reciprocity, assortment arises from information-mediated contingent behavior rather than population structure.

In the simplest reputation-based idealization, reputational labels track past cooperative behavior in expectation, and individuals condition their actions on observed reputation: when reputation is observed, cooperation is preferentially directed toward individuals labeled as “good,” whereas when reputation is unobserved, interaction is effectively random. Under this behavioral rule, conditioning on whether the focal individual’s reputation is observed yields

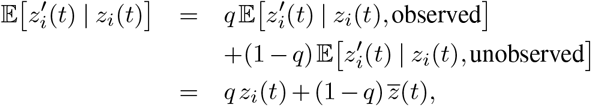

where 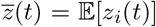 is the population mean cooperation rate. Because this dependence is linear in *z*_*i*_(*t*) with slope *q*, the resulting Price-equation regression coefficient of the partner’s cooperative response on the focal individual’s action reduces to *q* under this idealization, and therefore the effective assortment coefficient for indirect reciprocity is *r*_ind_ := *q*. Substituting *r*_ind_ into the Price-equation condition reproduces the indirect-reciprocity rule of (Nowak, 2006).

#### (iv) Network reciprocity

In network reciprocity, individuals occupy the vertices of a *k*-regular graph and interact only with their neighbors. We consider death–birth updating, in which a randomly chosen individual is replaced by a copy of one of its *k* neighbors. Under weak selection, the Price equation reduces the condition for cooperation to *rb > c*, where

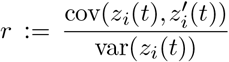

is the regression coefficient of a randomly sampled neighbor’s trait 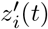 on the focal individual’s trait *z*_*i*_(*t*), evaluated under the neutral process. A simple genealogical heuristic explains why *r* scales as 1*/k*: tracing lineages backward under neutral death–birth updating, ancestral lineages perform a simple random walk on the graph; consequently, a randomly sampled neighbor coincides with the focal individual’s most recent ancestral step with probability of order 1*/k*, generating positive focal–neighbor correlation of that order. The exact value of *r*, however, is a derived property of the full neutral replacement chain and must be computed using pair-approximation or coalescent methods. For death–birth updating on large *k*-regular graphs, this calculation has been carried out by (Ohtsuki et al., 2006), yielding an effective assortment coefficient *r*_net_ = 1*/k* to leading order, which when substituted into the Price-equation condition recovers the network-reciprocity rule *b/c > k*.

#### (v) Group selection

Group selection is often formulated using a two-level Price decomposition, but for consistency with (Nowak, 2006) we work at the individual level, where the effects of group structure appear through induced covariance. Consider a population subdivided into *m* groups of size *n*, and let *z*_*i*_(*t*) ∈ {0, 1} denote the cooperative trait of individual *i* at time *t*. Writing 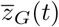 for the mean trait in group *G*, the law of total variance gives

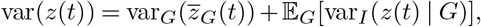

the standard decomposition into between- and within-group components. Under weak selection in the donation game, group-level competition implies that the relevant covariance term in the Price equation depends only on between-group differences,

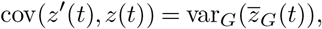

so that the assortment coefficient may be written as

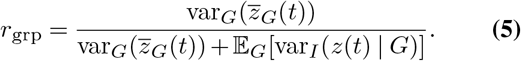

In the rare-splitting, weak-selection regime considered by (Nowak, 2006), the two-stage fixation-probability calculation of (Traulsen and Nowak, 2006) determines the effective neutral variance components induced by the group-splitting dynamics. In particular, for that model one obtains

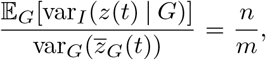

where *n* is the group size and *m* the number of groups. Substituting this ratio into Eq. (5) yields

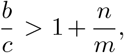

which is precisely the group-selection rule derived in (Nowak, 2006). This also shows that, for this class of models, group selection is mathematically equivalent to kin selection, with the assortment coefficient arising from between-group variance rather than genealogical relatedness (Okasha, 2006; Frank, 2013).

The three-step structure of this derivation reveals that the biological content of each rule resides entirely in step 2, i.e., the computation of the assortment coefficient *r* from the interaction or updating mechanism, while step 3, the Price equation, is identical across all five rules. The five mechanisms are therefore not five cooperation theories but five different ways of generating the same covariance structure.

### Transmission

Nowak’s five rules would be of interest, even as special cases of the Price equation, if they constituted a canonical and exhaustive list of mechanisms by which cooperation can increase. However, this is not the case. The Price equation reveals mechanisms that fall completely outside the scope of the five rules, because (Nowak, 2006) omits an entire component of the Price equation: the transmission term. Specifically, mutation-driven cooperation does not rely on assortment, repeated interaction, reputation, or group structure, and therefore cannot be reduced to any of Nowak’s five rules.

#### Corollary 2

Consider a population with a binary cooperation trait *z*_*i*_(*t*) ∈ {0, 1} . Suppose reproduction is neutral with respect to *z*, and that trait change occurs only through asymmetric transmission, with defectors producing cooperative offspring with probability *µ* and cooperators producing defective offspring with probability *ν* per reproduction event. Then the Price equation implies that the mean cooperation level increases at time *t* if, and only if,

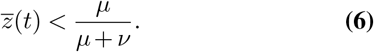

Equivalently, cooperation increases whenever its current frequency lies below the transmission balance determined solely by mutation.

*Proof:* Reproduction being neutral with respect to the trait means that cov(**w**(*t*), **z**(*t*)) = 0. Evolutionary change is therefore entirely driven by the transmission term. Under the two-way mutation model, the expected change in the mean trait due to transmission is

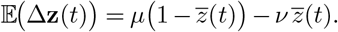

Under the usual neutral normalization in which 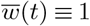, the Price equation reduces exactly to

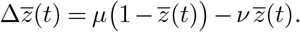

Rearranging yields

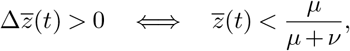

which is Eq. (6). Thus, in the absence of any selective covariance, cooperation can increase purely through biased transmission, independently of assortment, interaction structure, or payoff effects.

This result is, of course, classical (Fisher, 1930; Haldane, 1932; Price, 1970; Frank, 1997, 2012; Gardner, 2020), though typically formulated in terms of allele frequencies rather than cooperation as a trait. Hamilton’s rule characterizes when selection generates an instantaneous increase in cooperation due to population structure, whereas the transmission inequality instead describes a directional pull toward a transmission-defined equilibrium.

### Mechanism

The Price equation derivations above require only (i) a first-order (weak-selection) expansion of fitness and (ii) a specification of the joint law of a focal individual and its partner(s), summarized by an assortment coefficient. No particular evolutionary dynamics is required. Nevertheless, it is useful to exhibit a concrete microscopic model in which the assortment coefficients can be computed from first principles. A natural choice is a Moran process (Moran, 1958), because it is a well-posed continuous-time Markov chain on a finite state space (Ewens, 2004), whose dynamics are described exactly by a chemical master equation (CME). Moreover, under weak selection its drift admits a straightforward first-order expansion, making it easy to identify the mechanism-dependent assortment coefficient.

Fix a population size *N* and let **X**(*t*) = (*X*_1_(*t*),…, *X*_*T*_ (*t*)) ∈ ℕ^*T*^ denote the counts of *T* types at time *t*, with 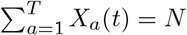. A continuous-time Moran process is a compositional continuous-time Markov chain on the finite state space, i.e.,

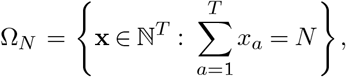

specified by a collection of reactions of the form

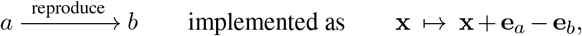

where **e**_*a*_ is the *a*th standard basis vector. Let *ρ*_*a*_(**x**) denote the reproduction propensity of type *a* in state **x** and let *κ*_*b|a*_(**x**) denote the probability that an offspring of *a* replaces an individual of type *b*. The resulting transition rate from **x** to **x** + **e**_*a*_ − **e**_*b*_ is

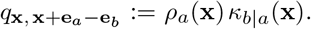

Writing *P* (**x**, *t*) for Pr(**X**(*t*) = **x**), the CME (Kolmogorov forward equation) is

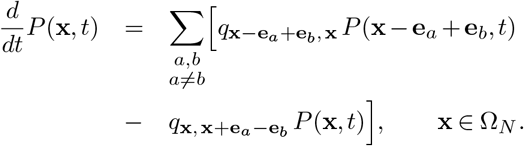

Equivalently, the generator *L* acting on test functions *f* : Ω_*N*_ → ℝ, which describes the evolution of expectations of observables, is

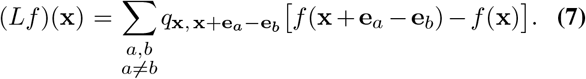

To connect the dynamics to cooperation, payoffs are introduced through a weak-selection perturbation of reproduction propensities. A standard parameterization is

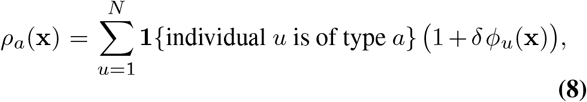

for 0 *< δ* ≪ 1 where *ϕ*_*u*_(**x**) is the (state-dependent) payoff-based contribution to fitness of individual *u* when the population configuration corresponds to **x**. The replacement kernel *κ*_*b*| *a*_(**x**) encodes the update rule and the interaction architecture: in a well-mixed birth–death Moran process one typically has *κ*_*b*| *a*_(**x**) = *x*_*b*_*/N*, whereas on a graph with death– birth updating, *κ*_*b* |*a*_(**x**) depends on local neighborhoods. Under Eq. (8), the generator admits the expansion

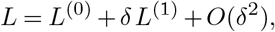

where *L*^(0)^ is the neutral replacement generator (obtained by setting *δ* = 0) and *L*^(1)^ is the first-order selection perturbation (Kimura, 1964). Thus, for any observable *f*,

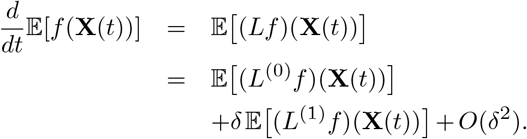

Taking *f* to be the mean cooperation level (i.e. 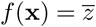 in the two-type case), the *O*(*δ*) term yields an explicit drift expression whose coefficients depend on the joint distribution of focal and partner types induced by *κ*_*b* |*a*_(·) and the interaction structure. This is precisely where the assortment coefficient enters: it is the scalar that summarizes the first-order correlation between focal and social environment generated by the neutral process together with the update rule.

In practice, computing the assortment coefficient reduces to computing neutral expectations under the generator *L*^(0)^. In structured populations, these expectations can be expressed in terms of meeting probabilities, coalescence times, or stationary solutions of linear systems associated with *L*^(0)^. The resulting coefficient depends on the mechanism: genealogical correlations in kin selection, temporal persistence of partners in direct reciprocity, information persistence in indirect reciprocity, spatial locality and update rule in network reciprocity, or group-level sampling and replacement in group selection. The advantage of the CME formulation is that each of these mechanisms corresponds to a precise choice of rate functions in Eq. (7), so that the assortment coefficient emerges as a derived property of the neutral process rather than an imposed parameter. Notably, there is no reason to restrict the CME framework to Moran processes. More generally, cooperation can be studied using CMEs that arise in other stochastic population models such as those used in molecular systems biology, where neutral generators and weak perturbations play an analogous role (Gorin et al., 2023).

Crucially, the cooperation inequalities themselves do not depend on Moran dynamics. Once weak selection yields a linear fitness perturbation and the induced focal–partner correlation is specified, the Price equation produces a cooperation inequality independently of the microscopic evolutionary process. Different population models may therefore give rise to identical first-order correlations and hence to the same cooperation conditions. For example, Wright–Fisher reproduction (Fisher, 1930), other continuous-time birth–death chains, or deterministic limits can all induce the same assortment coefficient and lead to identical cooperation inequalities (Frank, 1997). Moran processes underlie the derivations in (Nowak, 2006) because they provide an exact finite-population CME description, make the weak-selection expansion in Eq. (8) transparent, and offer a canonical route for computing assortment from interaction structure and update rules, not because they are essential to the inequalities themselves.

This separation of roles also clarifies a long-standing debate surrounding inclusive fitness. The criticism that inclusive fitness “fails” because it is not a dynamical theory (Nowak et al., 2010) rests on a category error. Inclusive fitness was never intended to generate evolutionary dynamics. Rather, it provides first-order conditions for selection once a population process has been specified. Its role is diagnostic rather than generative: given an underlying evolutionary model, inclusive fitness expresses the Price selection term cov(**w, z**) in a particular coordinate system, obtained via a regression decomposition of fitness effects when such a linearization is available. In this precise sense, inclusive-fitness reasoning is fully compatible with, and can be derived as a specialization of, the Price equation rather than constituting an independent dynamical theory.

Explicit population-genetic models such as Moran or Wright–Fisher processes are required to define evolutionary dynamics and to compute the coefficients that appear in the Price equation, including assortment or regression parameters. Inclusive-fitness expressions are posterior interpretations of these coefficients, not independent principles. When the structure of a model permits such a decomposition, inclusive fitness yields conditions equivalent to those obtained directly from the Price equation (Grafen, 1985; Frank, 1997; Grafen, 2006). When it does not, inclusive-fitness interpretations fail, but the Price equation itself remains valid. The apparent conflict between inclusive fitness and explicit evolutionary dynamics therefore reflects a confusion between identities that diagnose selection and models that generate population change.

Seen from this vantage point, the five rules for the evolution of cooperation (Nowak, 2006) are not alternatives to inclusive-fitness reasoning, but illustrations of it. Each rule corresponds to a particular mechanism by which population structure induces a correlation between the trait of a focal individual and the traits of its interaction partners. Once this focal–partner correlation is specified, the condition for cooperation follows from the Price equation under weak selection. The rules differ only in how assortment is generated, not in the underlying logic of selection. Their common structure reflects the fact that, to first order, cooperation inequalities are properties of selection under weak perturbations and are independent of the specific microscopic evolutionary dynamics used to compute the relevant correlations.

### Existence

The cooperation inequalities derived above rely on a weak-selection expansion of the discrete Price equation, under which only first-order fitness effects contribute. While this expansion is often presented as a Taylor approximation around neutral selection, it acquires a more natural interpretation after a change of coordinates. In particular, expressing fitness in additive rather than multiplicative form aligns weak selection directly with the underlying evolutionary dynamics. This change of coordinates is implemented by passing from Wrightian fitness to Malthusian fitness via a logarithmic transformation (Felce et al., 2025). In these coordinates, the continuous Price equation (2) replaces the discrete equation (1), and the associated deterministic dynamics take the form of replicator equations for relative frequencies.

Assuming faithful transmission, so that trait values are constant along lineages, the transmission term vanishes and the continuous Price equation reduces to a covariance identity. If **q**(*t*) ∈ Δ^*n*−1^ is the vector of type frequencies and **m**(**q**(*t*)) is the corresponding Malthusian fitness vector, then the replicator equation is

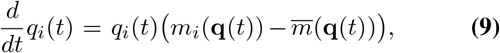

where

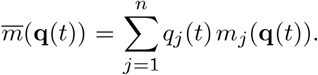

For any fixed trait vector **y** ∈ ℝ^*n*^, faithful transmission implies

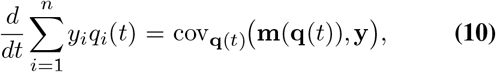

which is the continuous Price equation specialized to the absence of transmission bias.

As a motivating example, consider kin selection in the continuum. In this case there are two types (cooperator and defector), and individuals interact preferentially with genetically similar partners. Let *z*_*i*_ ∈ {0, 1} denote the cooperative trait, with *z*_*i*_ = 1 for cooperators and *z*_*i*_ = 0 for defectors, and let **z** = (*z*_1_, …, *z*_*n*_) be the corresponding trait vector, which is fixed under faithful transmission. Kin structure is encoded by a fixed interaction operator *G* = (*G*_*ij*_), where (*G***z**)_*i*_ represents the expected cooperative environment experienced by type *i*. We assume

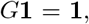

so that constant trait configurations are preserved.

Under weak selection, consider a donation game in which cooperators pay a cost *c >* 0 to their own Malthusian fitness and confer a benefit *b >* 0 to their interaction partners. To first order in the selection strength, the Malthusian fitness vector may be written as

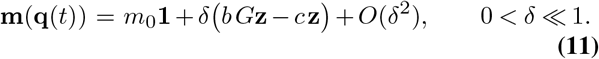

Here the full fitness may in principle depend on the current state **q**(*t*), but under weak selection we retain only the first-order term about the neutral reference dynamics.

Applying Eq. (10) with **y** = **z** yields

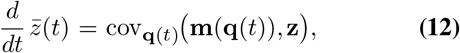

where

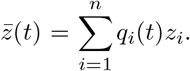

Substituting Eq. (11) into Eq. (12) and using the bilinearity of covariance together with

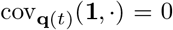

gives, to first order in *δ*,

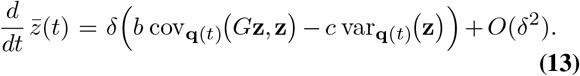

When var_**q**(*t*)_(**z**) *>* 0, define the relatedness coefficient

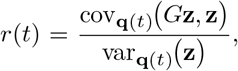

which measures the regression of the social environment on the focal trait at time *t*. Equation Eq. (13) may then be written as

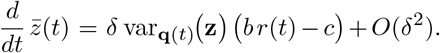

Thus, to first order in weak selection, cooperation increases locally in time whenever

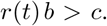

In this sense Hamilton’s rule emerges as the local growth condition for cooperation in the continuum limit.

This calculation suggests a more general question. In the kin-selection setting, the trait vector **z** is fixed and binary, and the first-order selection term reduces to a scalar multiple of var_**q**_(**z**), so the cooperation condition collapses to a single inequality *rb > c*. In more structured populations, however, *G* need not act as a scalar on all centered directions: different linear combinations of cooperation traits may be amplified at different rates. The natural question is therefore not whether a particular trait vector grows, but whether any centered perturbation of type frequencies is amplified by the interaction structure. We now show that this question has a clean spectral answer.

Let ***π*** ∈ int(Δ^*n*−1^) be a neutral reference distribution. We write

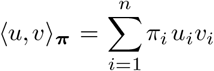

for the ***π***-weighted inner product, and

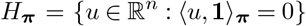

for the centered subspace. The point of introducing ***π*** is that, under weak selection, the first-order selection term is naturally evaluated at the neutral reference dynamics. In reversible settings, or more generally whenever the linearized interaction operator is self-adjoint in the ***π***-weighted geometry, the resulting first-order change is governed by a symmetric quadratic form.

#### Theorem 3

Let ***π*** ∈ int(Δ^*n*−1^) be a neutral reference distribution, and let *G* ∈ ℝ^*n*×*n*^ satisfy

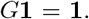

Assume that *G* is self-adjoint with respect to ⟨·, ·⟩_***π***_, and let *λ*_max_ denote the largest eigenvalue of the restriction 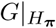.

Suppose that, to first order in weak selection and evaluated at the neutral reference distribution ***π***, the selection-induced change in the directional mean along **u** ∈ *H*_***π***_ is given by

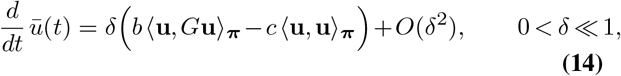

where *b, c >* 0 and

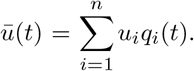

Then there exists a nonzero centered direction **u** ∈ *H*_***π***_ for which

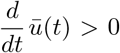

to first order in *δ* if and only if

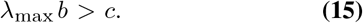

*Proof:* Because *G* is self-adjoint with respect to, ⟨· ·⟩_***π***_, its restriction 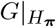 is also self-adjoint on the Hilbert space *H*_***π***_. Hence the Rayleigh quotient

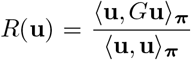

is well-defined for every nonzero **u** ∈ *H*_***π***_ and satisfies, by the Rayleigh–Ritz variational principle (Courant, 1920; Horn and Johnson, 2012; Carilli et al., 2025),

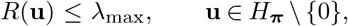

with equality for an eigenvector of 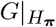 corresponding to *λ*_max_.

Now let **u** ∈ *H*_***π***_ be nonzero. Since

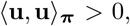

equation Eq. (14) implies that

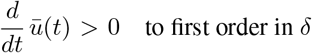

if and only if

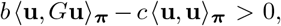

that is, if and only if

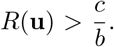

Therefore there exists a nonzero centered direction **u** ∈ *H*_***π***_ for which 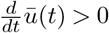 to first order if and only if

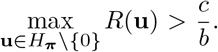

By the Rayleigh–Ritz variational principle,

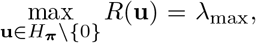

and hence this is equivalent to

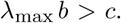

The spectral viewpoint reveals classical kin selection to be a degenerate case of this more general continuum framework, in which all centered cooperative directions are equivalent. Consider a population with neutral stationary distribution ***π***, and define the interaction operator

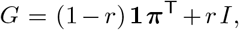

where *r* is the regression coefficient measuring assortment (relatedness) between interacting individuals. Acting on a trait vector **u** ∈ ℝ^*n*^, this operator yields

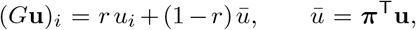

so that with probability *r* an individual samples its own trait, and with probability 1 − *r* it samples a random individual from the population. This is the standard mean-field representation of kin assortment.

Under weak selection, the first-order selection effect takes the form

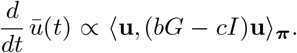

On *H*_***π***_, the rank-one operator **1 *π***^T^ vanishes identically, and hence *G* acts simply as

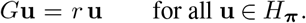

Thus every vector in the centered subspace is an eigenvector of *G* with eigenvalue *r*, and the leading eigenvalue relevant to the spectral condition is *λ*_max_ = *r*. The general spectral criterion *λ*_max_*b > c* therefore reduces exactly to *rb > c*, which is Hamilton’s rule (Hamilton, 1964).

In more structured settings, such as network, spatial, or reciprocity models, the interaction operator generally has a non-trivial spectrum on *H*_***π***_. Different linear combinations of individual cooperation traits are amplified at different rates. In such cases, no single scalar relatedness coefficient can summarize selection in all cooperative directions. The spectral criterion *λ*_max_*b > c* therefore provides a unifying and extensible condition for cooperation, capturing collective properties of the interaction structure that are invisible to any one-parameter reduction.

For example, consider a population of 3 types with uniform neutral stationary distribution

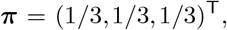

and suppose interactions are encoded by the reversible stochastic matrix

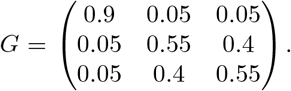

This matrix is symmetric and row-stochastic, so it is self-adjoint in the ***π***-weighted inner product and satisfies

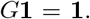

The constant vector **1** = (1, 1, 1)^T^ is therefore an eigenvector with eigenvalue 1, corresponding to the neutral mode eliminated by centering.

Restricting to the centered subspace *H*_***π***_, the remaining eigenvalues are 0.85 and 0.15. The leading centered eigen-vector is proportional to

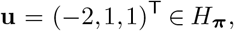

so the leading eigenvalue of *G* on *H*_***π***_ is

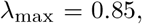

and the spectral cooperation condition is

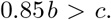

Choosing, for instance, *b* = 1.35 and *c* = 1, the leading mode is favored because

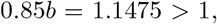

whereas the orthogonal centered mode with eigenvalue 0.15 is not, since

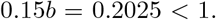

Thus the leading centered eigenvalue, rather than a single relatedness coefficient valid uniformly across all directions, is the relevant quantity in this structured setting.

This example illustrates the distinction between the two viewpoints. A scalar-relatedness description compresses the interaction structure to one number, whereas the spectral criterion *λ*_max_*b > c* asks whether there exists a centered cooperative mode, here the contrast between type 1 and the average of types 2 and 3, that is amplified by the global interaction structure.

## Discussion

We have shown that the five rules (inequalities) for evolution of cooperation (Nowak and May, 1992; Nowak and Sigmund, 1998; Traulsen and Nowak, 2006; Ohtsuki et al., 2006; Nowak, 2006) are a straightforward consequence of (Price, 1970) under weak selection. Notably, none of these works explicitly invoke the Price equation in their derivations, despite the fact that each rule can be recovered directly from the same covariance identity. The Price perspective also clarifies the parameters characterizing each cooperative mechanism in the payoff matrices in Table 1 of (Nowak, 2006). Those matrices should be understood as effective payoffs, obtained after averaging over interaction structure, repetition, or information flow so that standard replicator dynamics reproduces the desired cooperation condition. In our formulation, the payoff matrix encodes only the outcome of a single social interaction, namely the donation game in which cooperation confers a benefit *b* on the recipient at a cost *c* to the actor, while all mechanism-specific features enter through the covariance between an individual’s trait and the traits of its interaction partners. This viewpoint also subsumes later weak-selection results for structured populations, such as the *σ*-rule formulation (Tarnita et al., 2009, 2011), which can be understood as an alternative parameterization of the same Price equation covariance under particular update rules and population architectures. In the same vein, the payoff transformations introduced in (Taylor and Nowak, 2007) are also not fundamental objects in their own right, but artifacts of the weak-selection expansion of the Price equation. Treating them as primary obscures the fact that the same transformations arise generically whenever selection acts through covariances.

The Price equation perspective also forces a reckoning with the number of rules in (Nowak, 2006). First of all, Nowak’s five rules for the evolution of cooperation are evidently just a single selection rule. Second, the additional transmission rule can yield many more inequalities or “rules”. Notably, several prior studies have examined how mutation weakens cooperation in structured populations, most notably the analysis of evolutionary games on graphs with mutation (Allen et al., 2012). In that work, mutation enters the theory indirectly through its effect on identity-by-descent probabilities, which quantify spatial assortment under neutral drift and are computed using coalescing random walks. Cooperation conditions are then expressed as explicit benefit-to-cost thresholds in which mutation appears only via a reduced effective assortment coefficient. From the perspective developed here, these results correspond to a restricted special case of the present framework: mutation is not treated as a dynamical component of the evolutionary operator itself, but rather is absorbed into a single scalar summary of neutral correlations, after which selection is analyzed under weak-selection assumptions. By contrast, Eq. (6) arises directly at the level of the Price equation, where mutation contributes directly to the operator governing evolutionary change. This distinction clarifies both the scope and the limitations of earlier graph-based results, while showing how they fit naturally within a more general operator-theoretic formulation of mutation–selection balance.

Importantly, the transmission rule Eq. (6) is no more fundamental than the other rules. The unifying result on cooperation is the Price equation itself, which is not tied to any mechanism, model, or assumption, and is therefore the canonical organizing principle (Price, 1970; Queller, 1992, 2017). One criticism of the Price equation is that its universality stems from it being a tautology (Van Veelen et al., 2012; Nowak et al., 2017), and that models are necessary to derive insight into cooperation (Nowak and Highfield, 2011)^1^ But while the models in (Nowak, 2006) may constitute interesting toy examples with which to elucidate aspects of behavior, the inequalities of Table 1 are fundamentally a consequence not of the models but of the Price equation. Indeed, the Price equation makes clear that the suggestion in (Nowak, 2006) that “we might add ‘natural cooperation’ as a third fundamental principle of evolution beside mutation and natural selection” is futile, because the cooperation inequalities in (Nowak, 2006) derive from an equation involving only mutation and selection. That’s really all there is (Williams, 1965).

What is true is that the mathematics underlying the Price equation is elementary. The Price equation is essentially the product rule of calculus, yet it yields useful evolutionary insights in much the same way that the virial theorem yields insights in astronomy (Felce et al., 2025). The virial identity 2*K* + *U* = 0 expresses a balance between kinetic support and gravitational binding. When applied to perturbations of a homogeneous self-gravitating medium, it yields the Jeans criterion for instability, 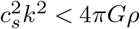, where *c*_*s*_ is the sound speed, *k* the perturbation wavenumber, *G* Newton’s gravitational constant, and *ρ* the mass density (Binney and Tremaine, 2011). In both cases, a general identity becomes predictive when coupled to system-specific structure, producing an inequality that governs stability or change. While the mathematics of the Price equation is elementary, its application is nontrivial. Confusion arises when these derived inequalities are mistaken for fundamental principles rather than specializations of a general identity, as emphasized by (Abbot et al., 2011; Rousset and Lion, 2011; Gardner et al., 2011; Boomsma et al., 2011; Strassmann et al., 2011; Bourrat et al., 2023). From this perspective, the Price equation provides insight at the level of principle (Gardner et al., 2011; Gardner, 2020; Lehtonen, 2020), whereas the specific Moran processes examined in (Nowak, 2006), while useful as illustrative models, do not themselves constitute fundamental explanations of cooperation. Treating such model-dependent inequalities as standalone principles risks mistaking illustration for explanation, a confusion that has also appeared in broader discussions of effective altruism (Burum et al., 2020). Evidently, despite the protestations of (Wilson, 2013), semi-literacy in mathematics is not sufficient for the study of eusociality (Frenkel, 2013).

We note that the spectral inequality we have derived, namely, *λ*_max_*b > c* (Eq. (15)), unifies a large body of work on evolution of cooperation. Importantly, it is derived in the setting of replicator dynamics, which play an important role in evolutionary theory. Replicator equations do not represent a specific microscopic population process but rather arise as deterministic limits of a broad class of stochastic evolutionary models under appropriate scaling regimes (Taylor and Jonker, 1978; Hofbauer and Sigmund, 1998; Page and Nowak, 2002; Rice, 2004). These include Moran and Wright–Fisher processes in large populations, continuous-time birth–death chains under weak selection, and chemical master equations describing population or molecular abundances. In this sense, replicator dynamics capture a broadly applicable first-order response of trait frequencies to selection, independently of the details of reproduction, updating, or demographic stochasticity.

Furthermore, replicator dynamics emerge naturally when fitness differences induce smooth changes in population composition and transmission is faithful. The resulting dynamics describe the projection of selection onto the space of trait frequencies, with the covariance structure of the population encoding all information about interaction patterns and population structure. This makes replicator dynamics the appropriate setting for identifying model-independent cooperation conditions under weak selection. The spectral formulation clarifies why this limit is so powerful. Once selection acts linearly on traits, the dynamics are governed by a linear operator acting on centered trait configurations, and the growth or decay of cooperation is determined by the leading eigenvalue of this operator. The inequality *λ*_max_*b > c* therefore plays the role of a linear stability condition for cooperation, analogous to instability criteria in other areas of applied mathematics (Strogatz, 2024). Importantly, this criterion depends only on the interaction operator and not on the microscopic details of the evolutionary process used to compute it.

The role of the Price equation and inclusive fitness are now also clear: the Price equation characterizes the instantaneous direction of evolutionary change by identifying the tangent vector to the trajectory of trait frequencies induced by selection. In this sense, it is a statement about the local geometry of the evolutionary flow on the simplex, regardless of how that flow is generated. Inclusive fitness corresponds to a particular coordinate representation of this tangent vector, obtained by projecting the selection gradient onto a chosen trait direction (Fig. 1). Neither object specifies a global trajectory in frequency space. Expecting the Price equation or inclusive fitness to generate evolutionary dynamics, as (Nowak et al., 2010) does, is therefore analogous to expecting a vector field to be a dynamical system. Conflating local geometric descriptors with generative dynamics mistakes the analysis of selection for a model of population change, and from a mathematical standpoint the debate over the status of inclusive fitness and the Price equation is settled by showing that they provide exact local descriptions of selection wherever a linearization of evolutionary dynamics is valid.

**Fig. 1.**
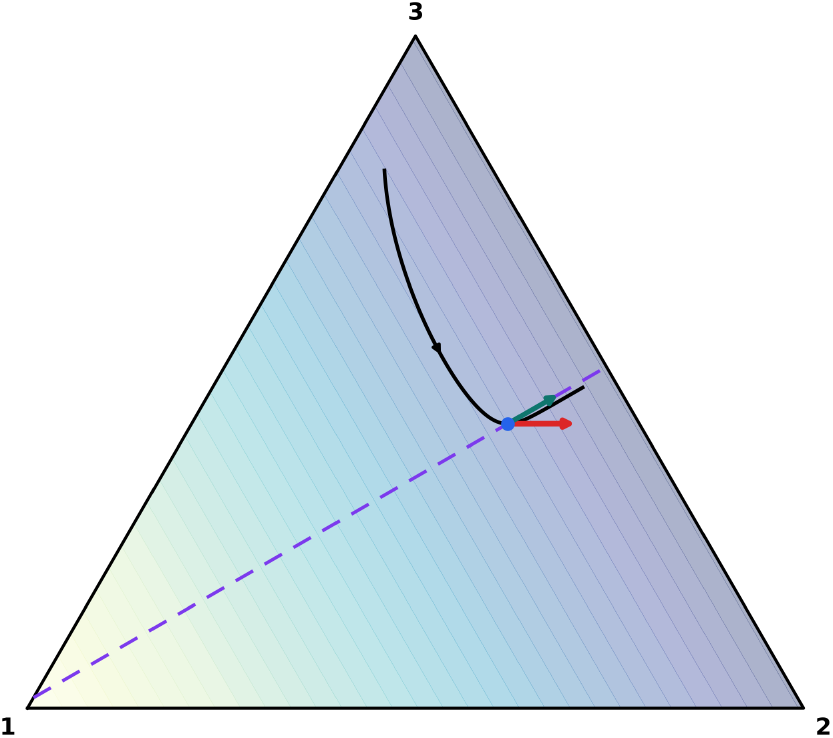
Geometry of weak selection in frequency space. The simplex (triangle) is the state space of relative type frequencies **q** = (*q*_1_, *q*_2_, *q*_3_) ∈ Δ^2^, for the reversible 3 × 3 example discussed in the text. The black curve is a trajectory **q**(*t*) generated by replicator dynamics Eq. (9), and the small black arrow indicates its direction of motion. The blue point marks a representative state **q**(*t*_0_) on this trajectory. The red arrow is the instantaneous Price direction 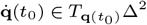, which satisfies 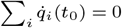. The background shading shows the cooperative mode identified by the leading centered eigenvector **u** ∝ (−2, 1, 1). The violet dashed line is the corresponding cooperation axis in the tangent space, and the teal arrow is the component of the Price direction along this axis. The red and teal arrows are drawn on the same display scale, so the teal arrow shows the projected contribution of the full Price direction to the cooperative mode. The code to reproduce this figure is available at https://github.com/pachterlab/P_2026/.

Moreover, this viewpoint unifies continuous and discrete formulations of cooperation. The classical rules derived for discrete-time or finite-population models correspond to evaluating the same spectral condition for finite-dimensional operators associated with specific update rules and interaction structures. Time discretization, population finiteness, or stochastic updating affect how the operator is constructed, but not the form of the cooperation condition itself. In this way, the spectral inequality provides a common language for results derived using Moran processes, Wright–Fisher models, graph-based dynamics, and their deterministic limits, revealing them as instances of a single underlying principle rather than distinct mechanisms. Furthermore, the spectral formulation immediately extends to settings in which the full structure of the operator matters. For heterogeneous networks (Chung, 1997; Allen et al., 2017), weighted interactions, or spatially extended populations (Geritz et al., 1998), the leading eigenvalue reflects collective properties of the interaction architecture rather than a single assortment coefficient. In such cases, cooperation thresholds cannot be reduced to scalar rules, and genuinely new predictions arise from the global geometry of interactions. This highlights the primary advantage of the spectral viewpoint: it provides a principled route to studying cooperation in complex populations for which no closed-form rule is available.

We emphasize that the present results concern local conditions for the increase of cooperation under weak selection. The spectral inequality *λ*_max_*b > c* governs the initial growth or decay of cooperative traits near neutrality and does not characterize global dynamics, fixation probabilities, or longterm equilibria under strong selection or nonlinear payoff structures. Extending the operator-theoretic framework beyond linear perturbations is an interesting direction for future work, for example by replacing the leading eigenvalue criterion with higher-order spectral quantities arising from tensor-valued interaction structures (Lim, 2005; Ribot et al., 2025) in systems with multi-way (Gokhale and Traulsen, 2014; Bick et al., 2023) or nonlinear interactions.

## Footnotes

1 The book *SuperCooperators: Altruism, Evolution, and Why We Need Each Other to Succeed* (Nowak and Highfield, 2011) quotes (Van Veelen et al., 2012), invoking a remark attributed to footballer Johan Cruyff and described as a tautology, namely that to win a soccer match one must score more goals than the opponent. This is used as an analogy for the Price equation: a statement that, while indisputably true, purportedly provides no insight into how to win. The analogy misfires for two reasons. First, Cruyff’s statement is not a tautology but a definitional truth, holding by virtue of the rules of the game rather than logical form. Second, while the Price equation does not prescribe a winning strategy, it does describe the instantaneous direction in which play is unfolding, i.e., who is advancing, who is retreating, and where pressure is building on the field. Inclusive fitness is then a particular way of expressing that local direction of play in chosen coordinates, e.g. along teammate-aligned axes. Dismissing such information as vacuous confuses useful live commentary on the flow of the match with a playbook for scoring goals, and mistakes a description of motion for an instruction on how to engineer it (Frank, 1995, 2012).

